# Transcriptional profiling of immune and inflammatory responses in the context of SARS-CoV-2 fungal superinfection in a human airway epithelial model

**DOI:** 10.1101/2020.05.19.103630

**Authors:** Claire Nicolas de Lamballerie, Andrés Pizzorno, Julien Fouret, Lea Szpiro, Blandine Padey, Julia Dubois, Thomas Julien, Aurélien Traversier, Victoria Dulière, Pauline Brun, Bruno Lina, Manuel Rosa-Calatrava, Olivier Terrier

## Abstract

Superinfections of bacterial/fungal origin are known to affect the course and severity of respiratory viral infections. An increasing number of evidence indicate a relatively high prevalence of superinfections associated with COVID-19, including invasive aspergillosis, but the underlying mechanisms remain to be characterized. In the present study, to better understand the biological impact of superinfection we sought to determine and compare the host transcriptional response to SARS-CoV-2 versus *Aspergillus* superinfection, using a model of reconstituted humain airway epithelium. Our analyses reveal that both simple infection and superinfection induce a strong deregulation of core components of innate immune and inflammatory responses, with a stronger response to superinfection in the bronchial epithelial model compared to its nasal counterpart. Our results also highlight unique transcriptional footprints of SARS-CoV-2 *Aspergillus* superinfection, such as an imbalanced type I/type III IFN, and an induction of several monocyte- and neutrophil associated chemokines, that could be useful for the understanding of *Aspergillus*-associated COVID-19 and but also management of severe forms of aspergillosis in this specific context.

## INTRODUCTION

The current pandemic of novel coronavirus disease 2019 (COVID-19), caused by severe acute respiratory syndrome coronavirus 2 (SARS-CoV-2) began in Wuhan, Hubei province, China, in December 2019. As of May 18, 2020, there have been more than 4,628,903 confirmed COVID-19 cases in the world as reported by the WHO, including 312,009 deaths (WHO). SARS-CoV-2 is a beta-coronavirus closely related to the severe acute respiratory syndrome coronavirus-1 (SARS-CoV-1) and the Middle East respiratory syndrome coronavirus (MERS-CoV) that emerged in 2003 and 2012, respectively. These viruses are also transmitted from animals to humans and cause severe respiratory diseases in afflicted individuals.

In a short period of time, significant effort has been devoted to understanding the molecular basis of the pathology associated with SARS-CoV-2 infection in an attempt to guide work on treatment, vaccine and diagnostic test development. Numerous clinical studies have reported the pathophysiology of COVID-19 has similar aspects to that initially described for SARS-CoV, *i.e*. acute lung injury due to over-inflammation following early stages driven by infection and viral replication (Chen et al., 2020; Guan et al., 2020; Huang et al., 2020; Mehta et al., 2020; Zhu et al., 2020). Nevertheless, the particular underlying mechanisms of these exuberant inflammatory responses in SARS-CoV-2 infection remain largely unknown and there is a need to expand our knowledge of the host’s response. In this context, several recent omics-based approaches, including *in vivo* and *in vitro* transcriptional profiling studies, have highlighted specific aspects of the signature of infection that could contribute to COVID-19 (Blanco-Melo et al., 2020; Gordon et al., 2020; Messina et al., 2020; Xiong et al., 2020). Blanco-Melo and colleagues, using transcriptional and serum profiling of COVID-19 patients, have notably shown that the SARS-CoV-2 infection signature was defined by low levels of Type I and III interferons juxtaposed to elevated chemokines and high expression of IL-6 (Blanco-Melo et al., 2020).

It is now well known that superinfections of bacterial/fungal origin can affect the course and severity of respiratory viral infections. For example, the co-pathogenesis of viruses and bacteria into the lung has been extensively studied, notably in the context of influenza superinfection by bacteria such as *S. pneumoniae* (Bosch et al., 2013; McCullers, 2014; Morens et al., 2008; Paget and Trottein, 2019). To date, there are limited data available on superinfections associated with COVID-19, though superinfections were reported in 10%-20% of SARS-CoV-2-infected adults admitted to Wuhan hospitals through the end of January 2020, and notably in 50%-100% of those who died (Zhou et al., 2020). In intensive care units, COVID-19 patients are at high risk of developing secondary infections, including fungal infections *e.g*. invasive pulmonary aspergillosis (Lescure et al., 2020). Indeed, a recent study on the French COVID-19 cohort reported that 33% of critically ill COVID-19 patients also showed invasive aspergillosis (Alanio et al., 2020). However, while the reasons for increased vulnerability to *Aspergillus* in COVID-19 patients remain undetermined, the putative contribution of *Aspergillus* to SARS-CoV-2 related lung inflammation and COVID-19 pathophysiology also constitutes a major unanswered question.

To better understand the biological impact of superinfection in the SARS-CoV-2 context we sought to determine and compare the host transcriptional response to SARS-CoV-2 versus that of a SARS-CoV-2 + *Aspergillus* superinfection. To reach this goal, we established a model of SARS-CoV-2 infection and superinfection in reconstituted human airway epithelia (HAE), based on previously published works (Nicolas de Lamballerie et al., 2019; Pizzorno et al., 2019, 2020). Our analysis reveals that both simple infection and superinfection induce a strong deregulation of core components of innate immune and inflammatory responses, however, it also highlights unique transcriptional footprints of the SARS-CoV-2 + Aspergillus superinfection that provide valuable insight for the understanding not only of *Aspergillus*-associated COVID-19 but also for the management of severe forms of aspergillosis.

## RESULTS

In order to identify similarities and differences between the host response to SARS-CoV-2 simple infection and *Aspergillus* superinfection, we sought to investigate the transcriptome of human respiratory epithelial cells during infection, in comparison with non-infected cells. Aware of the inherent limitations of experimental models using cell lines, we set up a model of infection/superinfection in a physiological model of reconstituted human airway epithelium (HAE). Developed from biopsies of nasal or bronchial cells differentiated in the air/liquid interphase, these HAE models, that we previously used with different respiratory viruses including SARS-CoV-2 (Nicolas de Lamballerie et al., 2019; Pizzorno et al., 2019, 2020) reproduce with high fidelity most of the main structural, functional and innate immune features of the human respiratory epithelium that play a central role during infection, hence constituting an interesting surrogate to study airway disease mechanisms. We infected nasal or bronchial HAE with SARS-CoV-2, and superinfection with *Aspergillus* was performed at 48 hours post-infection (hpi), which we had previously defined as the peak of acute SARS-CoV-2 infection in the HAE model (Pizzorno et al., 2020). Mock-infected, infected (CoV) and superinfected samples (CoV+Asp) were harvested at 72 hpi (24 h after superinfection) to perform mRNA-seq analysis (**Fig. 1A**). Both nasal and bronchial HAE models of superinfection were further characterized and validated in terms of viral production, impact on trans-epithelial resistance and apical release of IL-6, which we used as hallmarks of infection (**Fig. 2A**). Interestingly, in contrast with nasal HAE, we observed a significative increase on the relative viral production in the context of superinfection in bronchial HAE, associated with higher IL-6 levels and a stronger negative impact on trans-epithelial resistance (**Fig. 1B**).

**Figure 1.**
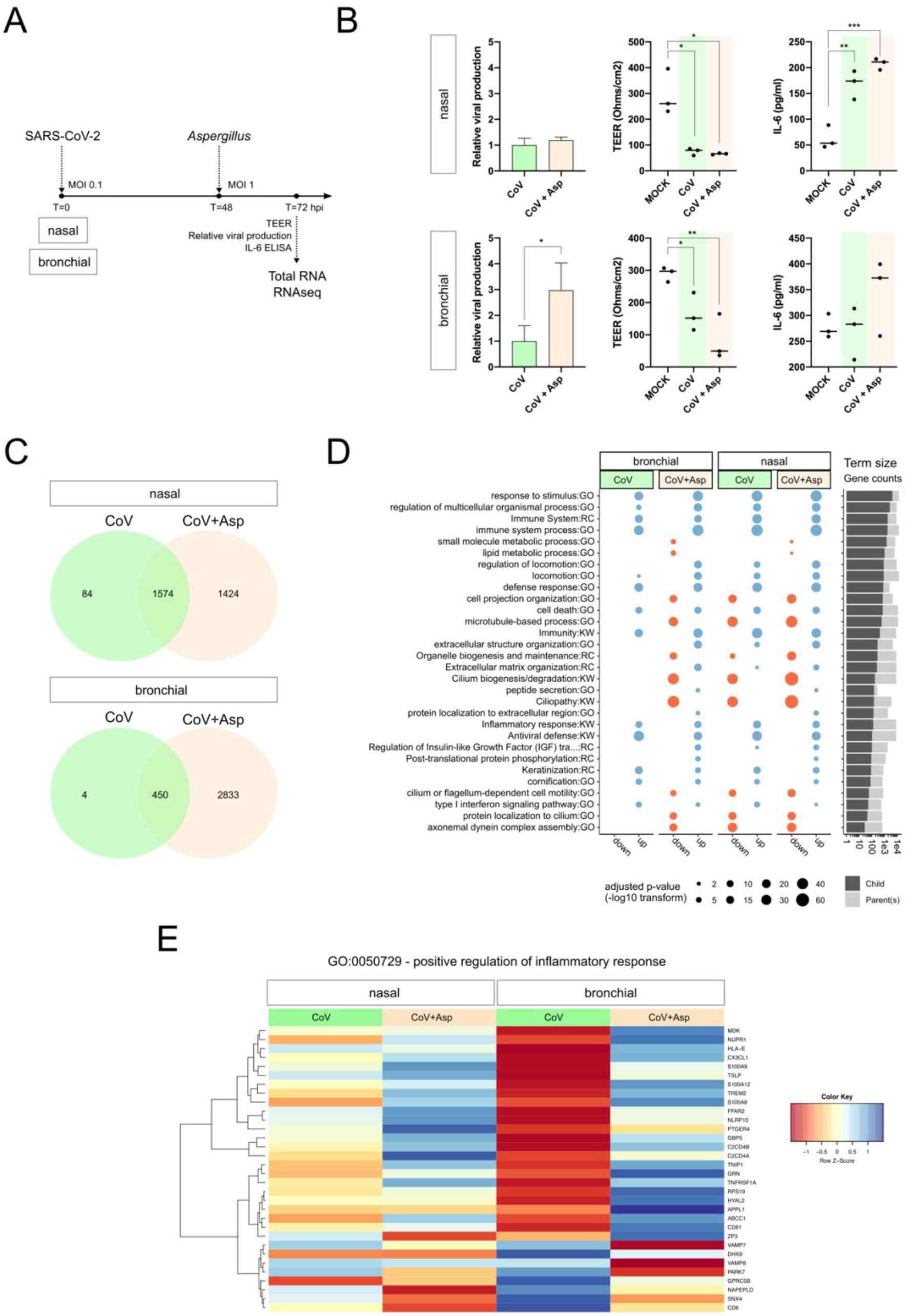
(A) Overview of experimental strategy (B) At 72h post-infection, for both nasal and bronchial HAE model, the relative viral production (intracellular) was determined using RTqPCR, and the impact of infection on epithelium integrity was monitored by measure of the transepithelial resistance (TERR Ohms/cm2). IL-6 was measured at the apical using a specific ALISA assay. (C) Nasal and bronchial gene signature overlap for SARS-CoV2 or SARS-CoV-2+Aspergillus infected conditions vs. Mock. We contrasted the significantly differentially expressed gene lists corresponding to CoV vs. Mock and CoV+Asp vs. Mock in nasal and bronchial HAE. Only genes above threshold (log2(FC)>1 or <-1 compared to the mock-infected condition and Benjamini-Hochberg adjusted p value < 0.01) were considered. (D) Overview of functional enrichment results for SARS-CoV2 or *SARS-CoV-2+Aspergillus* infected conditions vs. Mock. Considering (CoV vs. Mock) and (CoV+Asp vs. Mock) for both bronchial and nasal HAE, significantly up- or down-regulated gene lists (x-axis) were tested for significant enrichment using the parent-child strategy (see methods). If below the threshold (0.01), the adjusted p-values corresponding to different terms (y-axis) are represented by point sizes (see legend). After clustering terms based on gene occurrences (binary distance & Ward algorithm) in 15 metagroups, only the top 2 (lowest adjusted p-value) were represented here. For the complete list of significant terms, see supplementary figure S1. The bar plot on the right represents the sizes of enriched terms (called child) in comparison to the size of their parents (see methods for definitions). (E) Positive regulation of the inflammatory response in nasal and bronchial HAE. We extracted the list of proteins associated with the term GO:0050729 and visualized the scaled log transformed expression results in a heatmap. Each row represents a gene and each column is an experimental condition. The color key indicates the scaled expression levels vs. Mock (red, low; blue, high).

**Figure 2.**
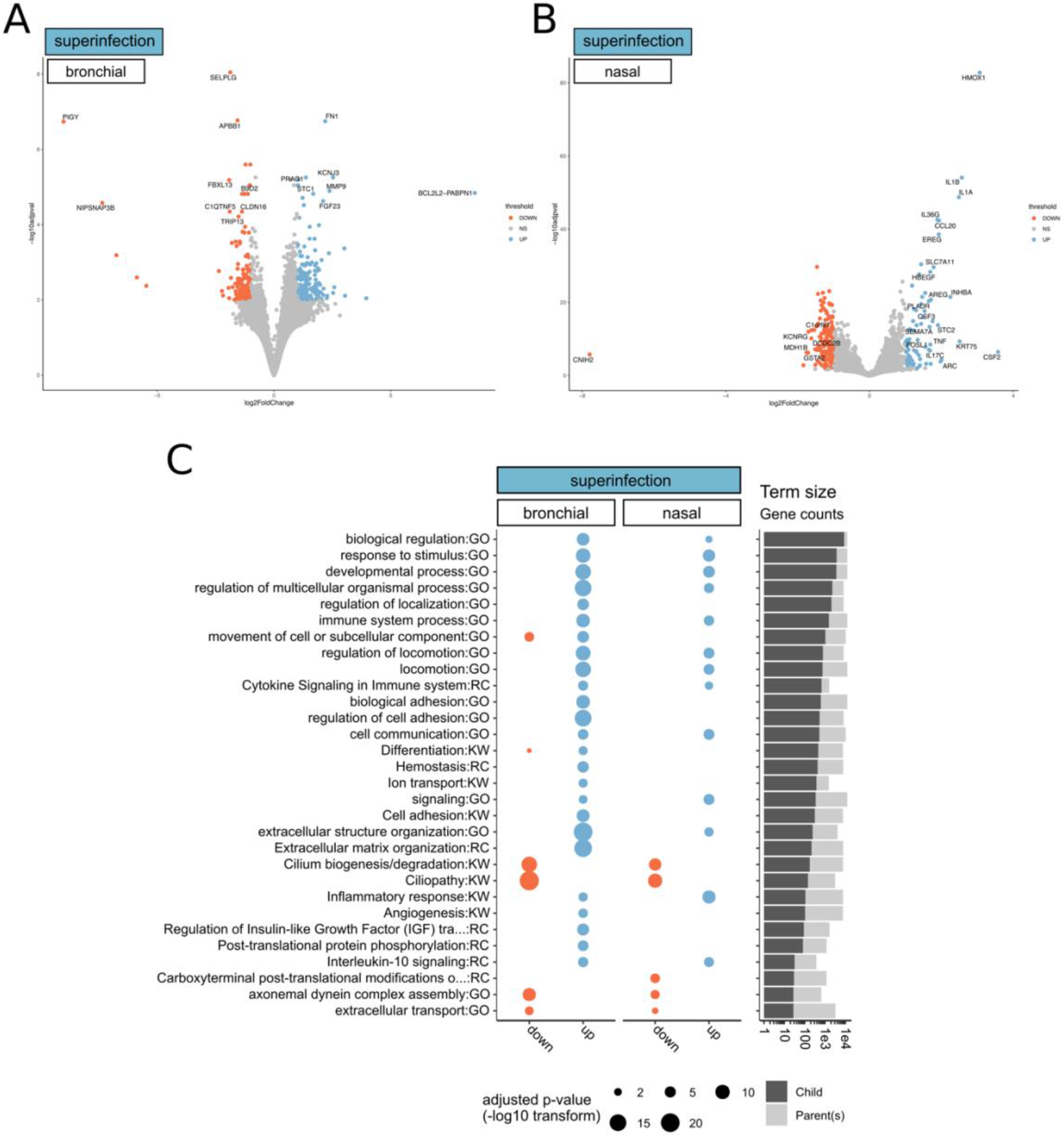
(A) & (B) Volcano plots of differentially expressed genes from (A) nasal and (B) bronchial tissue superinfected by *Aspergillus*. We considered significantly modulated genes (CoV+Asp vs. CoV) for both bronchial and nasal HAE (log2(FC)>1 or <-1 compared to the CoV condition and Benjamini-Hochberg adjusted p-value < 0.01). The color scale represents the log-transformed fold change values and ranges from low (red) to high (blue). (C) Overview of functional enrichment results of *SARS-CoV-2+Aspergillus* superinfection vs. SARS-CoV2 infection. Considering (CoV+Asp vs. CoV) for both bronchial and nasal epithelium type, significantly up- or down-regulated gene lists (x-axis) were tested for significant enrichment using the parent-child strategy (see methods). If below the threshold (0.01), the adjusted p-values corresponding to different terms (y-axis) are represented by point sizes (see legend). After clustering terms based on gene occurrences (binary distance & Ward algorithm) in 15 metagroups, the top 2 in terms lowest p-value were represented here. For the complete list of significant terms, see **Extended Data File 2**. The bar plot on the right represents the sizes of enriched terms (called child) in comparison to the size of their parents (see methods for definitions).

Differential expression analysis of mRNA-seq data compared to the mock-infected condition identified 1638 and 454 differentially expressed genes (DEGs) in SARS-CoV-2-infected nasal and bronchial HAE, respectively (FC ≥ 2, Benjamini-Hochberg adjusted p-value < 0.01). In the context of CoV+Asp superinfection, the number of DEGs was notably higher, with 2979 and 3235 genes in nasal and bronchial HAE, respectively (**Extended data. File 1**). Interestingly, there was an important overlap between CoV infection and CoV+Asp superinfection associated DEGs in both nasal and bronchial models. As illustrated in the Venn diagrams (**Fig. 1C**), more than 96% (nasal HAE) and 99% (bronchial HAE) of DEGs of the CoV signature were also part of the CoV+Asp signature. To provide further functional interpretation of the global transcriptional signatures, we performed a functional enrichment analysis on the CoV and CoV+Asp nasal and bronchial models using the web-based DAVID toolkit. Gene Ontology (GO), UniProt (KW) and Reactome (RC) terms where considered enriched when their Bonferroni-adjusted corrected enrichment p-value was < 0.01 (**Fig. 1D**). As anticipated, a large part of the most enriched and shared terms between all experimental conditions was related to the host response to infection (Regulation of response to cytokine stimulus, regulation of defense response, regulation of response to biotic stimulus) and also to cornification, which regroups genes mostly involved in cell-death mechanisms (**Fig. 1D**). Interestingly, functional enrichments specific to CoV infection or CoV+Asp superinfection were also highlighted. For example, CoV infection but no superinfection induced the upregulation of a gene cluster involved in Type I interferon pathway (**Fig. 1D**). Conversely, gene clusters harboring terms associated with epithelium physiology and cilium movements (regulation of cellular component movements, cell projection assembly, intraflagellar transport) but also with intra and extra-cellular signaling (post-translational protein phosphorylation, protein localization to extracellular membrane, peptide secretion) were exclusively enriched in the context of CoV+Asp superinfection (**Fig. 1D**). In parallel, given the importance of the exacerbated immune response observed mainly in severe COVID-19 cases, we further analyzed the inflammation-related terms and identified similar, yet to a different extent, regulation patterns for genes involved in the positive regulation of inflammatory response (GO:0050729) between nasal and bronchial HAE. As exemplified in the heatmap (**Fig. 1D**), deregulated genes such as calgranulin coding genes S100A12 and S100A9/A8 and downregulated genes such as Vesicle-associated membrane protein coding genes VAMP7 or VAMP8 were identified as a hallmark of CoV+Asp superinfection, in contrast to CoV infection (**Fig. 1D**). Similar observations were performed using different terms related to inflammation (**Extended data Fig. 1**), hence suggesting a very different inflammation signature resulting from superinfection. Altogether, our results indicate that the CoV+Asp superinfection presents a transcriptomic signature that recapitulates the overall signature of a simple CoV infection, but with both a particularly distinct regulation of the inflammatory response and the additional regulation of many biological processes related to the physiology of epithelia. Of note, we observed a relatively similar global pattern of regulation between the nasal and bronchial HAE models, differing primarily in the magnitude rather than the nature of the responses to CoV and CoV+Asp superinfection.

In order to explore in more depth the transcriptomic signature of the superinfection, we then performed a differential analysis of CoV+Asp signature using that of CoV infection as baseline. This focused analysis highlighted 248 and 769DEGs in the CoV+Asp nasal and bronchial HAE, respectively (FC ≥ 2, adjusted p-value < 0.01), which allowed us to spot finer differences between the two epithelium models. Indeed, the volcano plots in **Fig. 2A** and **B** representing all DEGs induced by the CoV+Asp versus the CoV condition show interesting differences in both the scale of deregulation and the nature of the most deregulated genes between the upper and lower respiratory tract tissues. Interestingly, many genes involved in the regulation of the inflammatory response such as HOX1, IL1B, IL1A, IL17C are found amongst the most upregulated genes in the nasal HAE model (**Fig. 2A**). These genes are also upregulated in the bronchial HAE model (**Extended data file 2**). To provide further functional interpretation of the superinfection signature, we performed a functional enrichment analysis, using the same strategy previously described. We only observed a limited number of enriched clusters of downregulated genes, mostly in bronchial HAE, all of them being related to epithelial physiology and cell movement (*e.g*. cytoskeleton-dependent intracellular transport, protein localization to cilium, cilium movement GO terms, **Fig. 2C**). In line with this observation, our analysis highlighted several clusters of enriched upregulated genes related to epithelial physiology (*e.g*. locomotion, movement of cell or subcellular component GO terms, **Fig. 2C**). On the other hand, terms related to signaling, host response and immunity were markedly more upregulated in the bronchial HAE model (**Fig. 2C** & **Extended data Fig. 2**). In addition to the gene enrichment profile shared between the nasal and bronchial models, our study also highlighted several overlapping biological processes such as the cytokine signaling and IL10 signaling pathways (**Fig. 2C**), which is consistent with the most upregulated DEGs shown in **Fig. 2A and Extended data file 2**. To better visualize these observations, we applied a protein-protein interactions analysis using STRING network to investigate the DEGs corresponding to several Reactome and GO terms (immune system process, cytokine signaling in immune system, inflammatory response, interleukin-10 signaling) enriched in the bronchial and nasal superinfection signatures along with their functional interactions (**Fig. 3A and 3B**). The two interactome maps with mostly upregulated DEGs illustrate the strong functional interconnections among several cytokines/chemokines (blue) and receptors (yellow) involved in the immune and inflammatory responses occurring in the context of a SARS-CoV-2 + *Aspergillus* superinfection. Interestingly, our analysis underlined the role of type III IFN (IFNL1, INFL2 and INFL3) and several cytokine-coding genes such as CXCL2/CXCL8 (**Fig. 3A** and **3B**) that are upregulated following simple SARS-CoV-2 infection and even more upregulated in the context of superinfection, arguably illustrating an enhanced specific response to control infection in both models.

**Figure 3.**
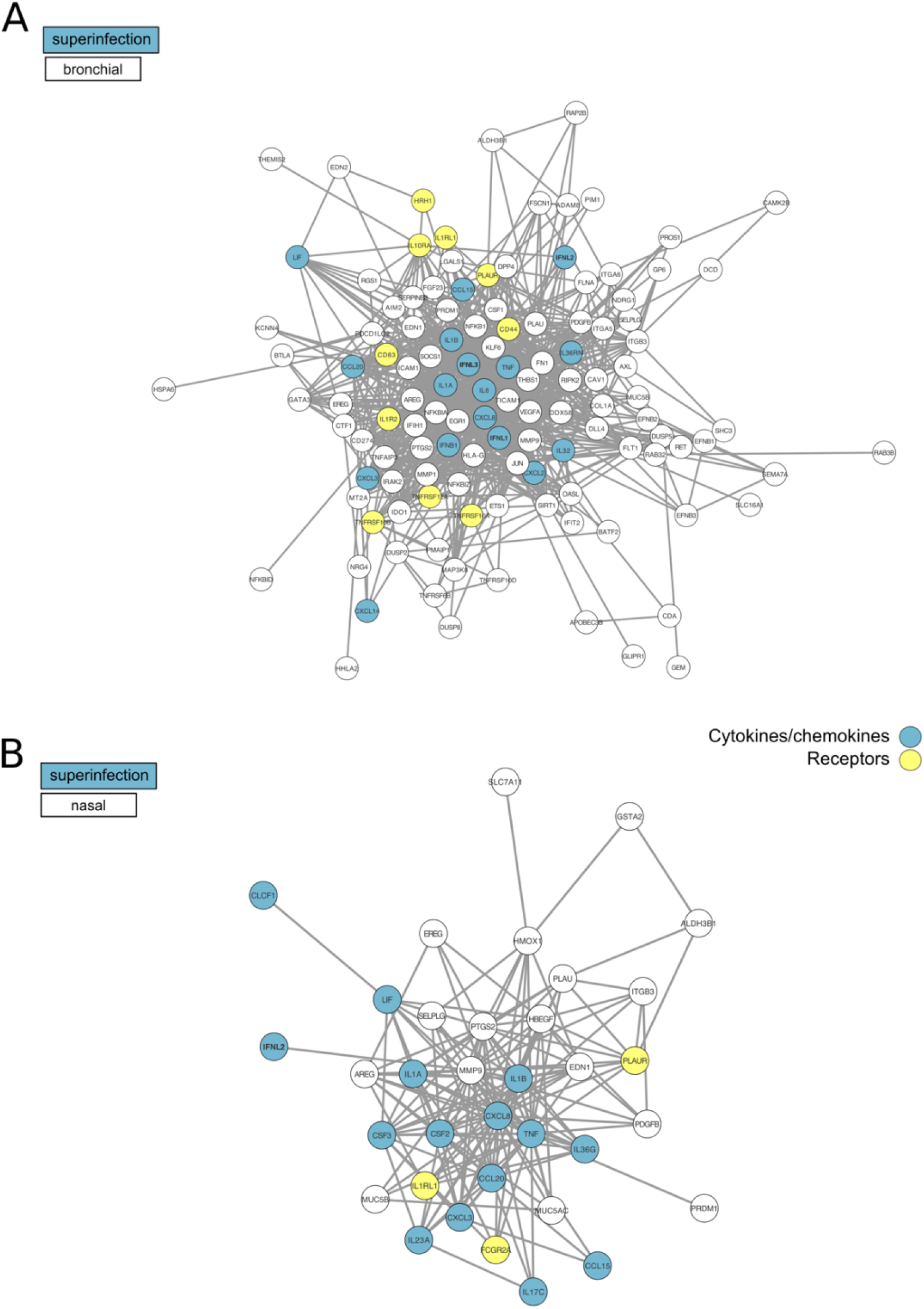
(A) & (B) Network visualization of immunity-associated proteins in superinfected HAE. We selected the 76 and 191 genes significantly differentially expressed (vs. CoV) in our study (respectively in nasal (A) and bronchial (B) HAE) associated with the Reactome “Interleukin-10 signaling” (R-HSA-6783783) and “cytokine signaling in immune system” pathway (R-HAS-1280215), the Uniprot keyword “inflammatory response” (KW-0395) and the Gene Ontology term “immune system process” (GO:0002376). The circles indicate genes modulated in our study in the superinfection context. The predicted associations are materialized by the dark lines. Cytokines/chemokines were highlighted in blue and receptors in yellow.

## DISCUSSION

Invasive pulmonary aspergillosis (IPA), which typically occurs in an immunocompromised host, represents an important cause of morbidity and mortality worldwide (Clancy and Nguyen, 2020). Superinfections were extensively documented in the case of influenza infections, with the latter being usually described to “pave the way” for bacterial superinfections, but several severe influenza cases have also been reported to develop invasive pulmonary aspergillosis 5/19/2020 9:59:00 AM. An increasing amount of evidence points towards a relatively high prevalence of superinfections, including invasive aspergillosis, to be associated with COVID-19 (Alanio et al., 2020; Lescure et al., 2020; Zhou et al., 2020). However, the underlying mechanisms remain to be characterized. In the present study, we sought to better understand the biological impact of superinfections by determining and comparing the host transcriptional response to SARS-CoV-2 versus SARS-CoV-2 + *Aspergillus* superinfection. Collectively, our results show a much stronger host response to superinfection in the bronchial epithelial model compared to its nasal counterpart. In both models, functional analyses show that the SARS-CoV-2 + *Aspergillus* superinfection signature reflects important changes in the expression regulation of genes involved not only in epithelium physiology but also in the regulation of host immune and inflammatory responses compared to that of the simple SARS-CoV-2 infection.

The reconstituted HAE model of infection/superinfection appears as a valuable support for the study of respiratory viral infections and virus-host interactions in highly biologically relevant experimental conditions. Previous results by our group using this model, constituted of fully differentiated and functional human primary cells, have provided meaningful contributions to the characterization of the kinetics of viral infection as well as on the tissue-level remodeling of the cellular ultrastructure and local innate immune responses induced by SARS-CoV-2, in line with our present observation (Pizzorno et al., 2020). Whereas no major differences in terms of global superinfection signatures where observed between HAE models of nasal or bronchial origin, the second part of our study highlighted more subtle differences between the two models in terms of scale of deregulation (fold change and p-value), as well as in the nature of the most deregulated genes (**Fig. 2C**). Ziegler and colleagues have recently reported that differences of infectivity and consecutive host responses between different cell subsets (type II pneumocytes, nasal goblet secretory cells) are linked to varying ACE2/TMPRSS2 levels, ACE2 expression being linked to the IFN response (Ziegler et al., 2020). The discrepancies we observed in the two HAE models could be explained by differences of cell type composition that could be interesting to further explore using combinations of additional experimental models, including ACE2/TMPRSS3 expression and single cell RNA-seq approaches.

Our analysis of the superinfection signature revealed an important role of physiology and cilium-related genes, which could reflect an additional negative impact of *Aspergillus* infection on the epithelium integrity (trans-epithelial resistance, **Fig. 1B**) and mucociliary clearance, in good agreement with previous observations by our group following different types of viral infection in HAE (Nicolas de Lamballerie et al., 2019). In the specific context of SARS-CoV-2 infection, the additional deleterious effect on epithelium integrity induced by aspergillosis might contribute to the enhanced disease severity reported in the clinic while arguably increasing the risk of additional bacterial of fungal superinfections, similarly to what has been described in the case of other respiratory viral infections (Smith et al., 2013; Wu et al., 2016).

Another major finding of our study relates to the impact of infection/superinfections on interferon and inflammatory responses, notably when compared to the limited transcriptomic data available in literature to date. Using primary cells, Blanco-Melo and colleagues recently reported that SARS-CoV-2 induces limited IFN-I and III responses (Blanco-Melo et al., 2020). Our data are not entirely consistent with these findings. Whereas we also demonstrate a very limited involvement of the type I interferon response in the context of simple SARS-CoV-2 infection -in contrast with data previously obtained with other viruses (Nicolas de Lamballerie et al., 2019), our analysis highlights an important upregulation of crucial genes involved in the type IIII interferon response. This response is even more pronounced in the context of superinfection (**Fig. 3A** and **3B**). Type III interferon (IFN-λ) is known to play a key role in innate and adaptive mucosal immune responses to infection (Ye et al., 2019). Interestingly, IFN-λ has been identified as a critical regulator of neutrophil activation to prevent fungal infection such as invasive pulmonary aspergillosis (Espinosa et al., 2017). Our data suggest a divergence between type I and type III interferon responses, with enhanced activation of the latter in the context of fungal superinfection. This characteristic response could be associated with increased neutrophil activation as an important first line of adaptive defense against these two pathogens. This hypothesis is further supported by the observed induction of several monocyte-and neutrophil associated chemokines, such as CCL2, CXCL2 and CXCL3 (**Fig. 3A** and **B**). Interestingly, our results suggest a strong activation of adaptive immune response in the context of SARS-CoV-2 infection, in line with currently available clinical data from COVID-19 patients, which generally present high levels of circulating neutrophils (Chen et al., 2020; Qin et al., 2020). These observations support an important role of neutrophil recruitment in the response to COVID-19, more particularly in the context of fungal superinfection with an exacerbation of pro-inflammatory response. In that sense, the upregulation of key genes belonging to the IL-10 pathway observed in our analysis (**Fig. 2C**), previously demonstrated to play a deleterious role in innate resistance to systemic aspergillosis (Clemons et al., 2000), constitutes an illustration of how such “enhanced” state of inflammation could contribute to increase severity.

In conclusion, our transcriptional profiling approach revealed unique features of the SARS-CoV-2 + *Aspergillus* superinfection signature, characterized on one side by an “enhanced” version of that induced by SARS-CoV infection, but also by specific changes on respiratory tissue physiology, a distinct regulation of type I and type III interferons, and an over-induction of inflammatory response. While we acknowledge that it would be rather bold to make a statement on the possible severity associated with *Aspergillus* superinfection in a pre-existing COVID-19 pathological context based solely on our results, our observations suggest that the immunomodulation induced by SARS-CoV-2 infection could establish a favorable context for the development of severe forms of aspergillosis. This could constitute an important aspect to be considered for the immunological follow-up of COVID-19 patients with aspergillosis. On top of that, the characteristic infection signatures described in this study provide valuable insight in the perspective of possible future treatments targeting the COVID-19 inflammatory response, which could result in counter-productive effects for the management of aspergillosis.

## MATERIALS AND METHODS

### Reconstituted human airway epithelial model

MucilAir™ HAE reconstituted from human primary cells obtained from nasal or bronchial biopsies, were provided by Epithelix SARL (Geneva, Switzerland) and maintained in air-liquid interphase with specific culture medium in Costar Transwell inserts (Corning, NY, USA) according to the manufacturer’s instructions. For infection experiments, apical poles were gently washed twice with warm OptiMEM medium (Gibco, ThermoFisher Scientific) and then infected directly with 150 μl dilution of virus in OptiMEM medium, at a multiplicity of infection (MOI) of 0.1 (Pizzorno et al., 2020). For mock infection, the same procedure was followed using OptiMEM as inoculum. Superinfection was performed in SARS-CoV-2 infected cells at 48h post-infection, with apical inoculation of 10 μl of *Aspergillus* in OptiMEM at a MOI of 1. Samples were collected from apical washes or basolateral medium at 72h post infection. Variations in transepithelial electrical resistance (TEER) were measured using a dedicated volt-ohm meter (EVOM2, Epithelial Volt/Ohm Meter for TEER) and expressed as Ohm/cm2.

### Pathogens

All experiments involving clinical samples and the manipulation of infectious SARS-CoV-2 were performed in biosafety level 3 (BSL-3) facilities, using appropriate protocols. The BetaCoV/France/IDF0571/2020 SARS-CoV-2 strain used in this study was isolated directly from a patient sample as described elsewhere (Pizzorno et al., 2020). Viral stocks were prepared and quantified in Vero E6 cells (TCID50/mL). *Aspergillus niger* (ATCC 16404) was quantified on maltose extract agar plates (CFU/ml).

### mRNA sequencing

Total RNA was extracted using the RNeasy Mini Kit (QIAGEN, ref 74104) with DNase treatment, following the manufacturer’s instruction. 500 μg total RNA was used to prepare polyA-enriched RNA-seq libraries using the KAPA mRNA Hyper Prep Kit (Roche, ref. KK8581/KK8581). Those libraries were prepared separately for each sample with 11 amplification cycles, then all libraries were equimolar-pooled before sequencing. Paired-end sequencing (2×100bp) was performed with Illumina NovaSeq 6000 sequencing platform on a SP flowcell (Illumina, ref: 20040326).

### RNA-Seq data trimming

Raw reads were first cleaned with Cutadapt 2.8 (Martin, 2011) to trim adapters (AGATCGGAAGAGCACACGTCTGAACTCCAGTCA and AGATCGGAAGAGCGTCGTGTAGGGAAAGAGTGT respectively for the first and the second reads) and low-quality ends (i.e terminal bases with phred quality score below 30). Only reads longer than 75bp and with less than 20% of N (undetermined base) after trimming were kept for further analysis.

### Transcript abundance quantification

The Kallisto 0.46.1 software was used for reference-indexing and transcript abundance estimation (Bray et al., 2016). The reference transcriptome based on NCBI RefSeq annotation release 109.20191205 and genome assembly build GRCh37.p13 were chosen for this analysis (Maglott et al., 2005). Default options were used for Kallisto except for reads orientation that was specified. The **Extended Data Table 2** provides a summary statistics of the pseudo-alignment process.

### Differential expression analysis

Differential expression analysis was performed with R 3.6.3. First abundances at transcript-level were imported using tximport 1.14.0 (Soneson et al., 2015). Only coding mRNA transcripts were considered. Gene-level effective lengths were obtained by weighted mean with their expression values in FPKM (Fragment Per Kilobase Million) weights. Raw counts were also computed at gene-level by sum and used as input for DESeq2 1.26.0 (Love et al., 2014). Normalization within DESeq2 used first gene-level effective length and then the size factor that was estimated using the function ‘estimateSizeFactors’. Differential expression testing was performed using default parameters. P-values were adjusted using the Benjamini-Hochberg method (Benjamini and Hochberg, 1995) after the independent filtering implemented by default in DESeq2. A gene was considered differentially expressed if both the adjusted p-value is below 0.01 and the induced change in expression is at least a two-fold increase (for up-regulated genes) or a two-fold decrease (for down-regulated genes). Finally, the NCBI gene IDs were mapped to Uniprot IDs using Uniprot cross-references (https://www.uniprot.org/database/DB-0118) (Breuza et al., 2016; Maglott et al., 2005). First, Uniprot entries associated to several gene IDs were removed. Indeed, we are not focused on gene fusion products or read-through transcripts in this study. In case of a gene ID associated to several Uniprot entries, only the best entry based on the reviewed status and the annotation score was kept; if those values were not discriminant enough, only duplicated entries with a reviewed status and an annotation score of 5 were kept. The protein-protein interaction (PPI) network was analyzed with STRING 11.0 and visualized with Cytoscape 3.8.0.

### In silico functional analysis

Based on mapped Uniprot IDs, gene set enrichment was performed using the parent-child union method as previously described (Grossmann et al., 2007). In this method each functional term, called child, is considered relatively to its parents to compute the probability of enrichment (p-value). We considered three functional databases with parent-child relations: Reactome (Fabregat et al., 2018), Uniprot Keyword (restricted to “Biological process” and “Disease”) (Breuza et al., 2016) and Gene ontology (Restricted to “Biological process”) (Ashburner et al., 2000; The Gene Ontology Consortium, 2015). Different files (available online: URLs were all accessed the 04/02/2020) were parsed to define Uniprot ID associations with terms and identify child to parents relations within each functional database; URLs are specified in **Extended Data Table 1**. As previously defined (Grossmann et al., 2007), the p_min probability represents how well a child term can be enriched relative to its parents. The higher this probability is, the less informative is the child term relative to its parents. This is also a statistic that can be used for independent filtering to reduce the p-value adjustment burden. Therefore, all terms with a p_min probability higher than 10e-4 were filtered out before p-values adjustment using Bonferroni method (Dunn, 1961). This multiple-testing correction procedure was handled separately for each functional database and analyzed gene list. A minimum adjusted p-value of 0.05 has been set for significant enrichment.

## Acknowledgments

The authors would like to thank Epithelix (Switzerland) and IGENSEQ sequencing core facility (Institut du Cerveau ICM, Paris) for their help. This study was funded by CNRS, and Mérieux research grants. The sponsors had no role in study design, collection, analysis and interpretation of data, manuscript writing, or in the decision to submit the article for publication.

## Extended Data

**Extended Data Figure 1.**
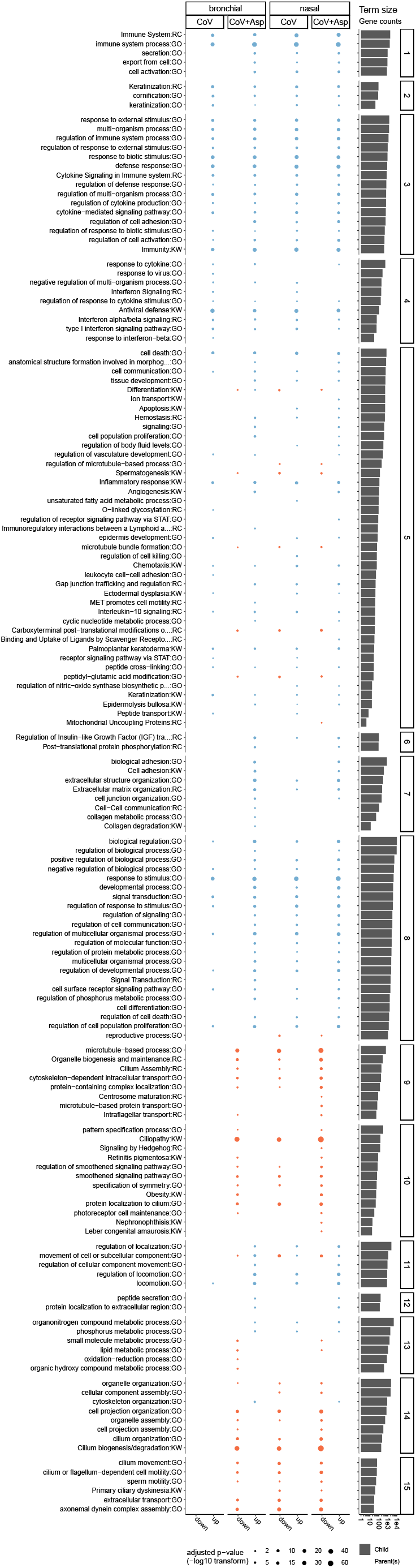
Complete view of functional enrichment results for SARS-CoV2 or SARS-CoV-2+Aspergillus infected conditions vs. Mock. Considering (CoV vs. Mock) and (CoV+Asp vs. Mock) for both bronchial and nasal epithelium type, significantly up- or down-regulated gene lists (x-axis) were tested for significant enrichment using the parent-child strategy (see methods). If below the threshold (0.01), the adjusted p-values corresponding to different terms (y-axis) are represented by point sizes (see legend). Terms were clustered based on gene occurrences (binary distance & Ward algorithm) in 15 metagroups. The bar plot on the right represents the sizes of enriched terms (called child) in comparison to the size of their parents (see methods for definitions).

**Extended Data Figure 2.**
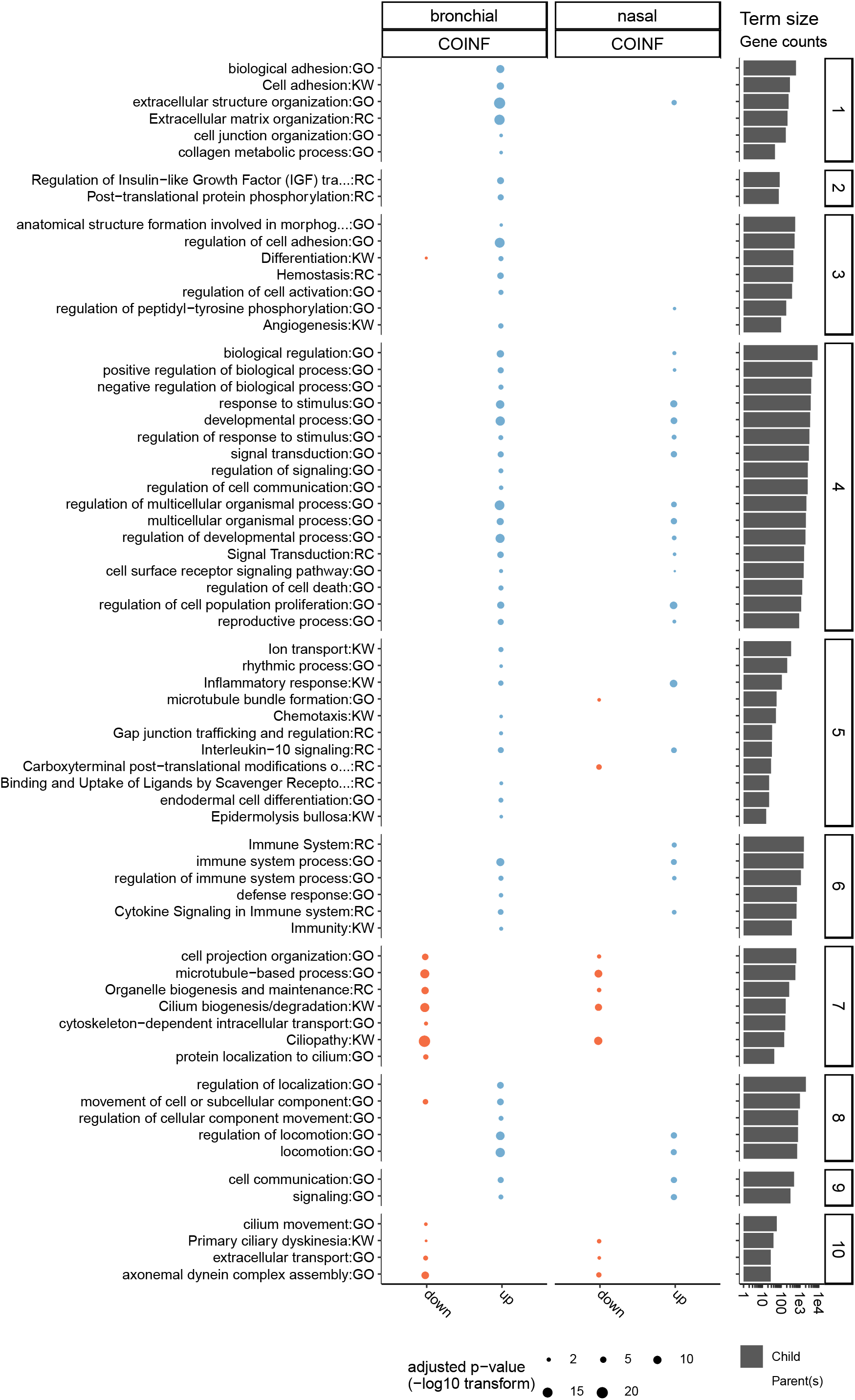
Complete view of functional enrichment results of *SARS-CoV-2+Aspergillus* superinfection vs. SARS-CoV2 infection. Considering (CoV+Asp vs. CoV) for both bronchial and nasal epithelium type, significantly up- or down-regulated gene lists (x-axis) were tested for significant enrichment using the parent-child strategy (see methods). If below the threshold (0.01), the adjusted p-values corresponding to different terms (y-axis) are represented by point sizes (see legend). Terms were clustered based on gene occurrences (binary distance & Ward algorithm) in 10 metagroups. The bar plot on the right represents the sizes of enriched terms (called child) in comparison to the size of their parents (see methods for definitions).

**Extended Data File 1 & 2**:

Extended data files will be available upon request

**Extended Data Table 1.** List of URLs parsed to build the databases used for enrichment analysis. Three types of information were used (i) details about terms meaning term ID and complete name and eventually category (e.g Biological process or Molecular function for Gene Ontology); (ii) gene-term associations; (iii) Topology and especially child to parent’s relations to apply the parent-child method.

| Database | Information type | URLs |
| --- | --- | --- |
| **Reactome** | Details (name,Id) | https://reactome.org/download/current/ReactomePathways.txt |
| **Reactome** | gene-term associations | [https://www.uniprot.org/uniprot/?sort=score&desc=&compress=no&query=organism:%22Homo%20sapiens%20[9606]%22&fil=&format=tab&force=yes&columns=id,entry%20name,length,genes(PREFERRED),reviewed,database(Reactome)](https://www.uniprot.org/uniprot/?sort=score&desc=&compress=no&query=organism:%22Homo%20sapiens%20[9606]%22&fil=&format=tab&force=yes&columns=id,entry%20name,length,genes%28PREFERRED%29,reviewed,database%28Reactome%29) |
| **Reactome** | Topology | https://reactome.org/download/current/ReactomePathwaysRelation.txt |
| **Uniprot Keywords** | Topology | http://ftp.ebi.ac.uk/pub/databases/interpro/ParentChildTreeFile.txt |
| **Uniprot Keywords** | Details (name,Id) | http://ftp.ebi.ac.uk/pub/databases/interpro/entry.list |
| **Uniprot Keywords** | gene-term associations | [https://www.uniprot.org/uniprot/?sort=score&desc=&compress=yes&query=organism:%22Homo%20sapiens%20[9606]%22&fil=&format=tab&force=yes&columns=id,entry%20name,length,database(InterPro),genes(PREFERRED),reviewed](https://www.uniprot.org/uniprot/?sort=score&desc=&compress=yes&query=organism:%22Homo%20sapiens%20[9606]%22&fil=&format=tab&force=yes&columns=id,entry%20name,length,database%28InterPro%29,genes%28PREFERRED%29,reviewed) |
| **Gene Ontology** | Topology and Details (name,Id) | https://www.uniprot.org/keywords/?sort=&desc=&compress=yes&query=&fil=&format=obo&force=yes' > keywords/uniprot_keywords.obo.gz |
| **Gene Ontology** | gene-term associations | [https://www.uniprot.org/uniprot/?query=organism%3A%22Homo%20sapiens%20%5B9606%5D%22&sort=score&columns=id%2Centry%20name%2Ckeywords%2Cgenes(PREFERRED)%2Cdatabase(GeneID)%2Creviewed&format=tab](https://www.uniprot.org/uniprot/?query=organism%3A%22Homo%20sapiens%20%5B9606%5D%22&sort=score&columns=id%2Centry%20name%2Ckeywords%2Cgenes%28PREFERRED%29%2Cdatabase%28GeneID%29%2Creviewed&format=tab) |

**Extended Data Table 2.** Statistics of RNA-Seq fragment pseudo-alignment to the human transcriptome.

| Epithelium type | sample | Percentage of fragments aligned | Millions of fragments aligned |
| --- | --- | --- | --- |
| Bronchial | CoV+Asp | 22,30% | 9,6 |
|  | CoV+Asp | 25,60% | 5,7 |
|  | CoV+Asp | 25,20% | 7,1 |
|  | CoV | 61,50% | 13,8 |
|  | CoV | 66,30% | 15,9 |
|  | CoV | 46,00% | 11,5 |
|  | Mock | 80,30% | 19,1 |
|  | Mock | 78,60% | 20,9 |
|  | Mock | 79,20% | 18,5 |
| Nasal | CoV+Asp | 45,40% | 10,8 |
|  | CoV+Asp | 47,00% | 15,1 |
|  | CoV+Asp | 40,90% | 9,5 |
|  | CoV | 53,00% | 15,2 |
|  | CoV | 53,00% | 14,4 |
|  | CoV | 57,10% | 15,7 |
|  | Mock | 82,20% | 21,7 |
|  | Mock | 80,70% | 22,1 |
|  | Mock | 80,40% | 19,2 |

## Notes

### Competing Interest Statement

The authors have declared no competing interest.

